# Signature of N-terminal domain (NTD) structural re-orientation in NPC1 for proper alignment of cholesterol transport: Molecular dynamics study with mutation

**DOI:** 10.1101/2020.06.09.141630

**Authors:** Hye-Jin Yoon, Hyunah Jeong, Hyung Ho Lee, Soonmin Jang

## Abstract

The lysosomal membrane protein NPC1 (Niemann-Pick type C1) and NPC2 (Niemann-Pick type C2) are main players of cholesterol control in lysosome and it is known that mutation on these proteins leads to cholesterol trafficking related disease, called Niemann-Pick disease type C (NPC) disease. The mutation R518W or R518Q on NPC1 is one of such disease-related mutations, causing reduced cholesterol transport by half, resulting in accumulation of cholesterol and lipids in late endosomal/lysosomal region of the cell. Even though there has been significant progress in understanding cholesterol transport by NPC1 in combination with NPC2, especially after the structural determination of full length NPC1 in 2016, many details such as interaction of full length NPC1 with NPC2, molecular motions responsible for cholesterol transport during and after this interaction, and structure and function relations of many mutations are still not well understood.

We report the extensive molecular dynamics simulations to gain insight into the structure and motions of NPC1 lumenal domain for cholesterol transport and disease behind the mutation (R518W). It is found that the mutation induces structural shift of NTD (N-terminal domain), toward the loop region in MLD (middle lumenal domain), which is believed to play central role in interaction with NPC2 protein, such that the interaction with NPC2 protein might be less favorable compare to wild NPC1. Also, the simulation indicates the possible re-orientation of the NTD, aligning to form an internal tunnel, after receiving the cholesterol from NPC2 with wild NPC1 unlike the mutated one, a possible pose for further action in cholesterol trafficking. We believe the current study can provide better understanding on the cholesterol transport by NPC1, especially the role of NTD of NPC1, in combination with NPC2 interaction.

**Synopsis:** modeling study of cholesterol binding protein NPC1

## 1. Introduction

The cholesterol homeostasis is maintained by many different proteins depending on tissues on the body.[1] Blood cholesterol levels are regulated by several processes, including bio-synthesis, cholesterol absorption/re-absorption, and biliary clearance and excretion.[2,3] Elevated blood cholesterol levels contribute to atherosclerotic coronary heart disease.[4,5] Previous studies have shown that lowering the levels of plasma cholesterol significantly reduces the risk of cardiovascular diseases associated with diabetes mellitus even in diabetic patients with normal levels of plasma cholesterol.[6] Therefore, control of cholesterol for therapeutic purpose for disease such as cardiovascular disease has fundamental importance. There has been tremendous amount of efforts to understand how different organ systems such as not only the liver and the intestine but also the brain, which are less explored, coordinate their cellular mechanisms to control body cholesterol homeostasis.[1,6]

The transmembrane protein Niemann-Pick type C1 (NPC1) inside lysosome is one of the key players in cholesterol transport, which is mediated by Niemann-Pick type C2 (NPC2).[7,8] NPC1 facilitate transport of LDL-derived cholesterol out of lysosomes for subsequent delivery to the endoplasmic reticulum and plasma membrane. NPC1 functions mostly in tandem with NPC2, a soluble lysosomal protein, to move unesterified cholesterol.[9] The abnormal function of NPC1 due to some factors such as mutation cause accumulation of cholesterol within lysosome, leading to a disease called NPC disease. Not only the crystal structures of *N*-terminal lumenal domain (NTD) of NPC1 with and without cholesterol (PDB id: 3GKI and 3GKH),[9] but also the cryo-EM (electron microscope) of full length NPC1 with and without cholesterol in NTD (PDB id: 3JD8 and 5JNX) were reported.[10] The NPC1 glycoprotein has 13 helixes in transmembrane domains (TMD) and three relatively large, lumenally oriented domains. The N-terminal lumenal domain (NTD) contains a cholesterol binding pocket while the newly determined structure of cysteine-rich C-terminal domain (CTD) contains loop region, which is close to the NTD, indicating the importance of this region in cholesterol transport in conjunction with maintaining the possible orientation of NTD to receive cholesterol from NPC2.[11] It is believed that the soluble NPC2 accepts LDL-cholesterol and delivers it to NTD directly, so called “hands-off mechanism”.[12] It is known that the two protruding MLD loops in NPC1 mediates interaction of NPC2 as well as interaction of glycoprotein of Ebola virus and both the structure of MLD with Ebola glycoprotein and NPC2 has been determined.[10,13-16] It facts, study of NPC1 has importance not only as a mediator of cholesterol transport but also as a mediator of coronavirus such as SARS and Ebola virus. Especially, much attention is payed to the NPC1 as a one of the possible target protein to mimic the NPC disease in relation to the recent pandemic outbreak COVID-19 virus because inhibition of this protein could reduce the replication of coronavirus including COVID-19 virus.[17,18]

There have been several computational or modeling studies regarding the cholesterol transport through NPC1, complementing the experimental results. The putative structure of NPC2 with cholesterol in complex with NTD was suggested with modeling studies.[19-21] The detailed sliding like hands-off cholesterol transport, especially the isomerization of cholesterol during the transport from NPC2 to NTD, was reported with QM/MM study within fixed NPC2/NTD conformational framework.[20] Based on the x-ray structure of NPC2 in complex with NPC1-MLD region, another possible NTD/NPC2 complex was also suggested.[16] The interface contacts between NTD and NPC2 are quite different between these two putative complexes, especially in terms of the angle between NTD and NPC2 and the orientation of NPC2 relative to the NTD. From the molecular dynamics simulations it is found that one of the complex, termed as “Texas model”, has favorable interaction in NPC2/NTD complex interface when the cholesterol is in NPC2 side while the same complex dissociates when the complex is in NTD side or no cholesterol on both side.[19,21] On the other hands, the other complex, coined as “California model”, shows the opposite behaviors. Based on these observations it was suggested that the Texas model may corresponds to the initial structure for cholesterol transport from NPC2 to NTD while the California model corresponds to the structure after the cholesterol transport.[21]

Even though there has been significant progress of understanding the dietary cholesterol exchange in cellular environment in connection to the NPC1 especially after the structural determination of full length NPC1, many of the details still remains unclear. The location or orientation of NTD presented in full length NPC1 structure might not in active form and there could have change of NTD orientation/location from full length NPC1 structure during the cholesterol transfer from NPC2 for proper alignment.[10,11,22] Eventually, it is believed that once the cholesterol has been transferred to NTD from NPC2 it should be delivered to the sterol sensing domain (SSD), which is in between the helix bundle within the membrane region, possibly triggering a sequence of events once transferred there.[23,24] The transfer of cholesterol to SSD could proceed from NTD possibly through a conduit like channel that was observed in Patched protein by proton driven network [25,26]. Note that the modeling study with disease causing L472P mutation indicate break down of this tunnel in NPC1,[27] emphasizing importance of this tunnel in cholesterol transport. Recent structure of NPC1 with NPC1 blocker itraconazole obtained from cryo-EM indicated the blocker is located in this tunnel, supporting this scheme.[28] In this context, the possible binding sites are suggested through modeling study recently.[29] We note that the channel was observed in short molecular dynamics simulation with no cholesterol in NTD.[30] Interestingly, the very recent molecular dynamics study NPC1 with cholesterol in itraconazole binding site indicates migration of cholesterol to SSD side along this tunnel with wild type unlike mutation, which migrates reverse direction.[30] Another possibility is that the cholesterol is transferred from NTD of neighboring NPC1, i.e. inter-protein transfer.[31] In either case, there should be some structural re-arrangement of NPC1 protein after the cholesterol is transferred to NTD from NPC2, possibly significant amount of NTD re-orientation or translocation for further actions.[16,29] Recent modeling study shows that the presence of cholesterol on SSD induces large structural change. i.e. unfolding of transmembrane 3(TM3), break of contact between MLD and CTD, and disengagement of NTD, suggesting that cholesterol on SSD can serve as either positive or negative feedback.[23]

The mutational study, both theoretically and experimentally, may provide opportunities to gain insight into the mechanism of NPC1 cholesterol transfer process directly or indirectly. Since the first discovery of mutation on NPC1 protein and its connection to NPC related diseases[32], numerous mutations either on NPC1 or NPC2 are additionally found.[11,33] The point mutation R518W (or R518Q) is one of such examples[32] and it has been reported that the cholesterol transfer activity is reduced by 50 % by this mutation.[10] At the same time, it is found that the binding affinity of NPC2 to the full length NPC1 is noticeably reduced.[10,12] The transfer efficiency was maximized under pH 5.5, which is the environment of lysosome.[12,34] From experiments, it has been reported that the interaction between NPC2 and NPC1 has two aspects, i.e. cholesterol independent weak interactions and cholesterol depending strong interactions.[12] Later, it has been found that this cholesterol dependent binding affinity is due to the structural difference of NPC2 near the MLD loops binding region depending on the presence of cholesterol.[16] There might be initial binding of NPC2 to NPC1 MLD loops and stable complex formation of NPC2 with NTD when there is cholesterol on NPC2, thereby the NTD is acting as an anchoring player also as pointed out by Gong et al,.[10] The computational study of point mutation I1061T, P1007A, and G992W on CTD, shows structural instability of the mutated NPC1, especially instabilities in NTD, suggesting that importance of correct orientation or stability of NTD in NPC1.[35] On the other hands, the computer simulation with NTD protein and its two mutations (Q92R and Q92S) shows the importance of correct electrostatic distribution near the entrance of cholesterol pocket as well as structural stability for suitable binding cholesterol to NTD.[36] In combination with experiment, the molecular dynamics simulation of L472P mutation indicates disruption of the tunnel between MLD and CTD that is believed to play essential role for cholesterol transport as mentioned earlier.[27] Since it was pointed that the R518W mutation decrease cholesterol transfer activity not because of misfolding of NPC1 but because of the functional defection of NPC1,[12] the atomic level detailed mechanism behind this defective functionality could provide insights on the structure and functional relationship of NPC1, especially its interaction with NPC2.

In this paper, we present extensive molecular dynamics simulations of NPC1 to understand the structural and dynamical characteristics upon R518W mutation and its effects on overall cholesterol transport efficiency in connection with possible re-location or orientation of NTD and interaction of NPC1 with NPC2.[32]

## Material and methods

We build the initial full length NPC1 structure using X-ray structure (PDB ID: 5U74) as a template.[11] The missing NTD structure was introduced by overlapping the overall structure into cryo-EM structure presented by Gong et al,, (PDB ID 3JD8).[10] For simulation with R518W mutation, we mutated the Arg518 to Trp518 with PyMOL[37] by selecting the lowest steric hindrance conformer. The pH of the simulation was set to 5.5 considering the lumenal environment of lysosome.[12,29,34] For this purpose, considering the intrinsic pKa values of amino acids, the protonation states of every Asp, Glu, and His was obtained by PROPKA 3.1[38] The overall structure was solvated with TIP3P water and the total charge was balanced by addition of Na^+^ ions. In this study, we only considered the lumenal exposed part of the full length NPC1 to reduce the overall computational cost by imposing position restraints on C_α_ atoms connected to TM. The restraints constant was set to 1000kJ/mol·nm^2^ and its locations are Pro259, Lys392, Glu610, Tyr871, and Tyr1088. Note that we have included the Proline-rich long strand that is connected to the NTD starting from membrane surface in our simulation. Of course, this simplified model may not fully reflect the behavior of full length NPC1 including membrane. The recent molecular dynamics simulation[30] with full NPC1 when cholesterol is present in NPC1 inhibiting itraconazole binding site,[28] which is at the interface between membrane and lumenal region, shows there is non-negligible distance correlation coefficient between TMD and the rest of the domains. However, the longtime simulation of full NPC1 suggest that the conformational stability of TMD is higher, i.e. not much structural change starting from initial structure with C_α_ RMSD below 2 Å, unlike the lumenal exposed three domains.[27] Also, one of the current simulation result is in agreement with the very recent full length NPC1 simulation, which will be addressed in Results section. Therefore, the model presented here might capture the qualitative structural/dynamic features of lumenal domains within simplified framework. The initial structure we have used in this study is shown in Figure 1. All figures of protein structure are drawn with PyMOL.[37] We have used CHARMM36 force field[39] for both protein and cholesterol when needed. The simulation box was constructed by setting the minimum distance from the protein edge to the simulation box as 10.0 Å and the final box size was 102.58, 110.22, and 83.82 Å along the x, y, and z-direction respectively. The periodic boundary condition was imposed along the three different directions. The non-bonded cutoff distance was set to 12.0 Å with pair list update at every 25 time steps. The long range electrostatic interaction was treated with particle-mesh-Ewald (PME) method with PME order of 4 and Fourier spacing of 1.2 Å. The initially prepared system was energy minimized with steepest descent method followed by molecular dynamics simulation with constraints to all-bond in protein using LINCS algorithm for 1 ns before the production run. For final production, the simulation was run under the constant temperature and constant pressure condition, i.e. NPT simulation. The temperature was set to 310K using velocity re-scale method with coupling constant of 0.1 ps and the pressure was set to 1 atm using Berendsen pressure coupling method with coupling constant of 2.0 ps with compressibility of 4.5×10^−5^ bar^-1^. The whole simulation was run using Gromacs-2018.6.[40] We used the leap-frog time integrator with time step of 4.0 fs to speed up the simulations by increasing the hydrogen atom mass by factors of four while keeping the total atomic mass unchanged by subtracting the same mass from the H-bonded heavy atoms as implemented in Gromacs.[41,42] We have performed the molecular dynamics for four different systems, i.e. wild NPC1, R518W mutated NPC1, wild NPC1 with cholesterol on NTD, and R518W mutated NPC1 with cholesterol on NTD. The reason we included system with cholesterol in NTD for mutated NPC1 in our simulation is we would like to trace the effect of mutation after successful cholesterol transfer from NPC2 to NTD since as much as half of cholesterol is transferred to NPC1 from NPC2 according to experiment in mutation.[10] The simulation time and total number of independent simulations are described in detail in Results section.

**Figure 1.**
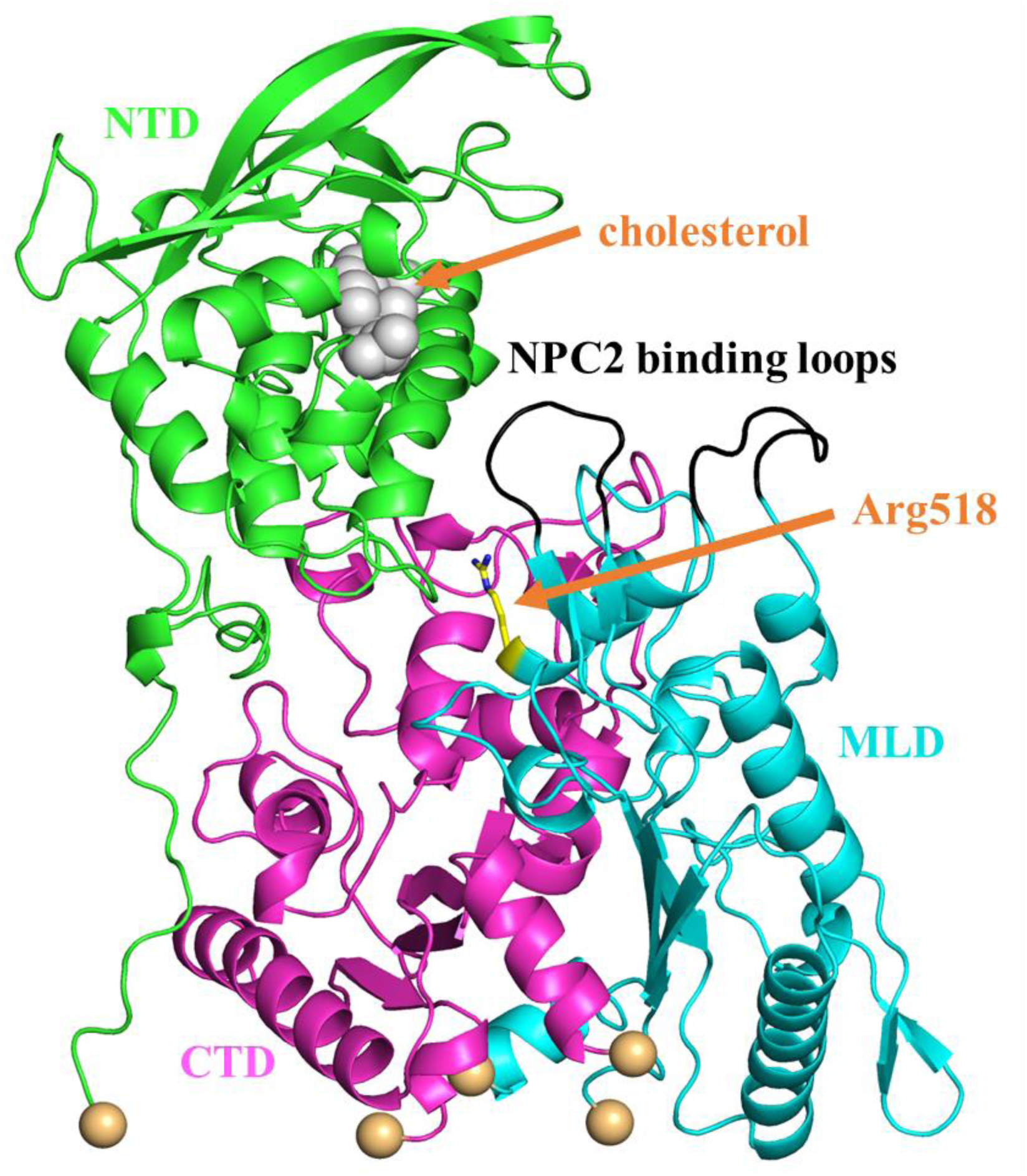
The initial structure we have prepared for simulations including cholesterol in NTD, which is shown as gray sphere. The NTD, CTD, and MLD domains are shown as green, magenta, and cyan respectively. The yellow stick is the Arg518 residue which is subject to the mutation to Trp in our simulation. The NPC2 binding loops are shown in black and the restrained C_α_ atoms are shown in wheat.

## Results and discussion

Initially, we run eight independent wild NPC1 simulations with no cholesterol in NTD for 200 ns. One of the simulation trajectories shows noticeable change of NTD displacement toward the outside of the NPC1 with RMSD of 8.7 Å compare to the initial structure. Therefore, with this trajectory we further proceeded the simulation up to 1.0 μs. The rest of the trajectories essentially show no noticeable change from the starting structure, exhibiting RMSD around 6.0 Å from starting structure. Among those remaining seven trajectories, we have randomly selected two more trajectories and proceeded the simulation to 1.0 μs but these two trajectories remains essentially steady. The results presented here are obtained from the trajectory with the large NTD displacement mentioned above. To monitor this displacement or tilting of NTD, we obtained the relative angles between each domains as a function of simulation time and showed in Figure 2. The angles shown here are relative angles between long helices from each domain. Note that the RMSD of each domain is relatively small during this simulations (less than 3.5 Å). Therefore, monitoring the angles between well preserved helices from each domain could be a measure of relative change of orientation between them. For this purpose, we defined a single vector from each helix and calculated the angles between them.[43] The helices we defined are shown in Supplementary Figure S1. Panel A shows sudden change of angles, especially angle between CTD and NTD, starting from around 100 ns and remains steady with some fluctuations after 300 ns. Compare to panel A, the same plot on panel B and C, which correspond to the angles in those two randomly selected trajectories, shows minor angle change. The Figure 3 shows the overlap of final structure from our 1μs simulation trajectory over the initial structure. One can clearly see the displacement of NTD away from the MLD domain. The movie file corresponding to this trajectory can be found in Supplementary material. In fact, this observation is in agreement with the simulation of full length NPC1 including the membrane with no cholesterol in NTD when the cholesterol was not present within the membrane.[23]

**Figure 2.**
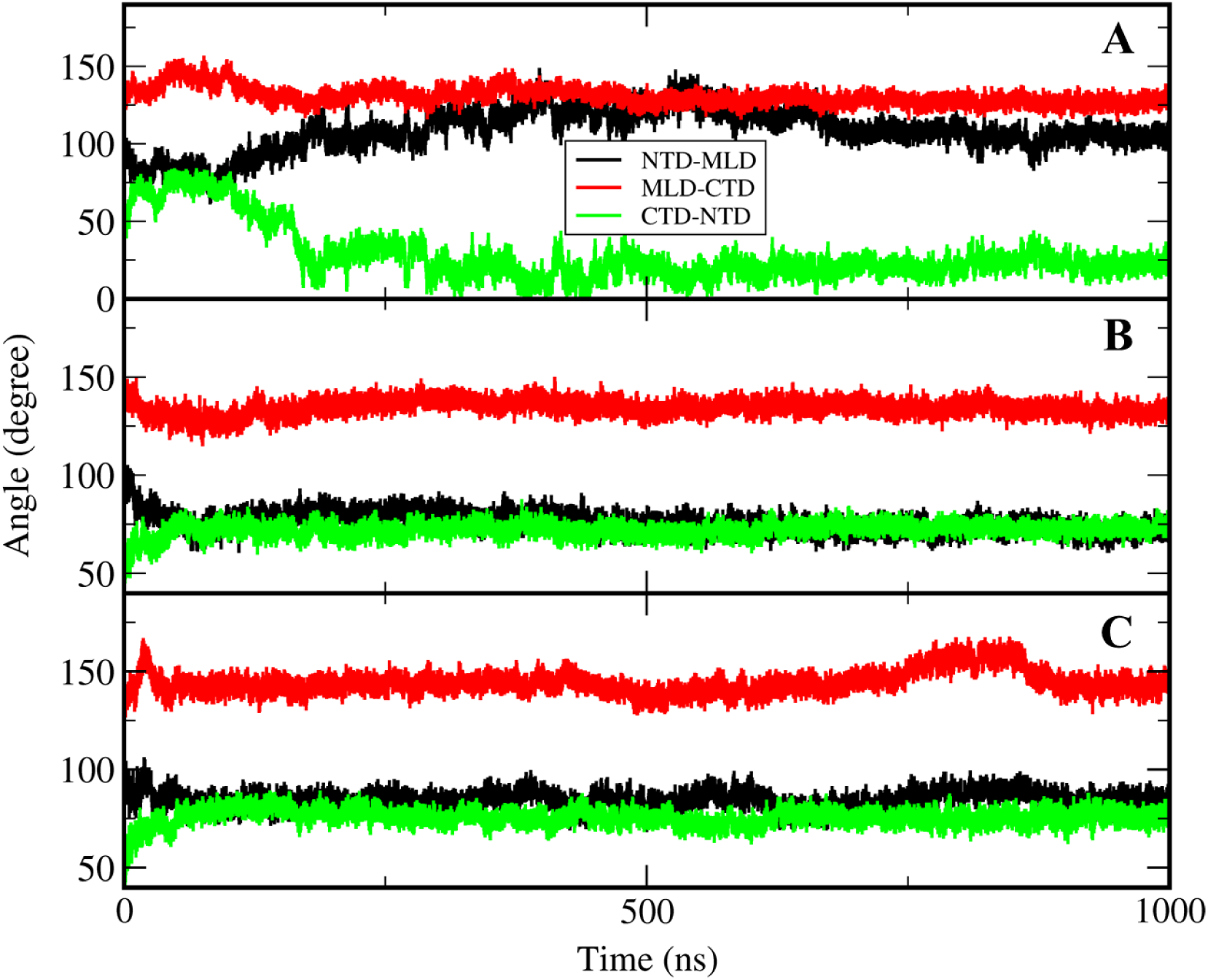
The time evolution of angles between NTD, MLD, and CTD for three trajectories. The panel A corresponds to the trajectory with titled NTD. We selected a helix from each domain and calculated the angles between each helix axis. Please refer Figure S1 for the selected helices in each domain.

**Figure 3.**
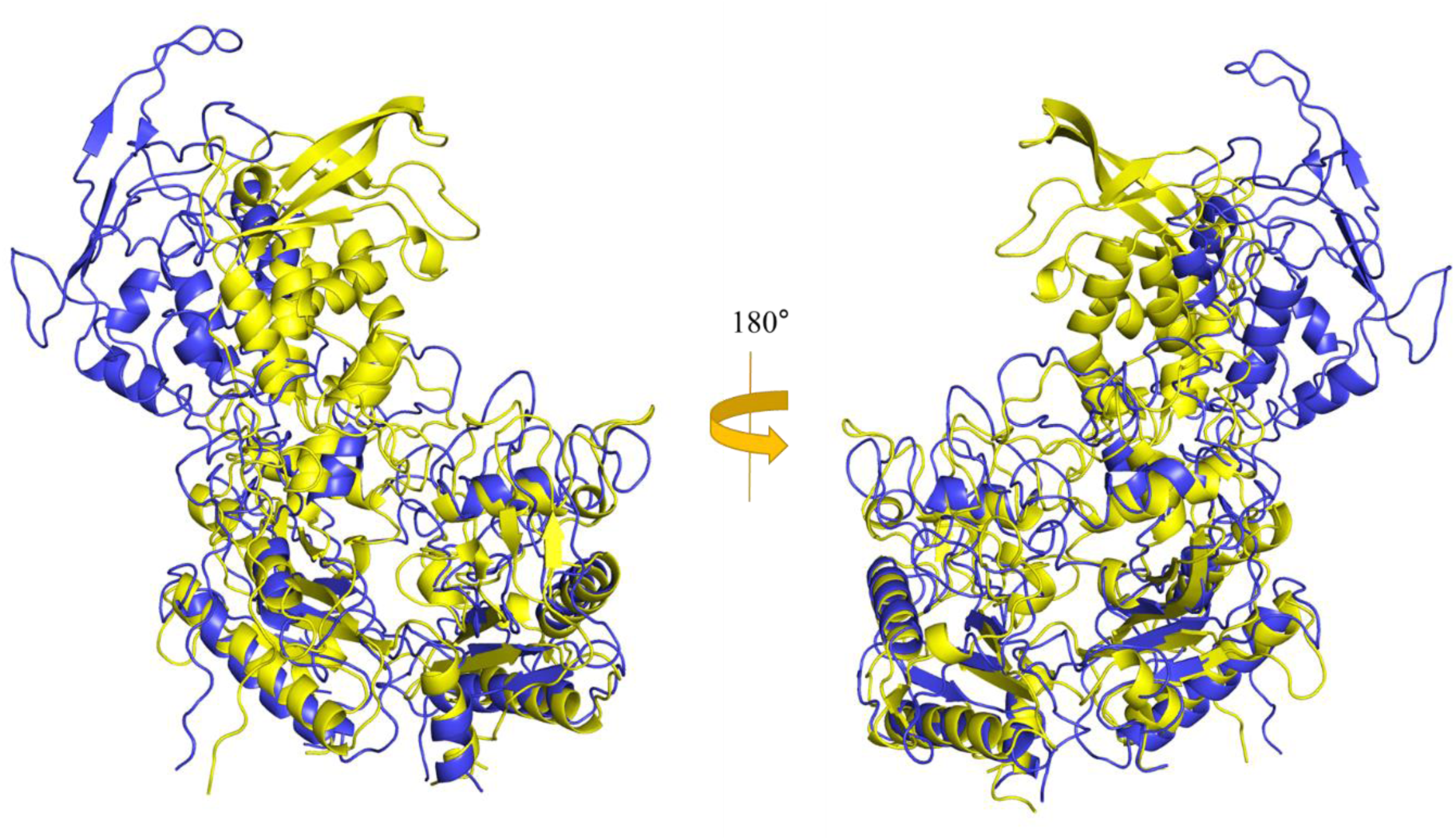
The structure obtained from current simulation with wild type after 1.0 μs (blue) that is in overlap with the initial structure (yellow).

The detailed structural information on the interaction or complex formation of NPC2 with NTD in the presence of full length NPC1 could significantly enhance our understanding on cholesterol transfer from NPC2 to NTD. We note that the overlap of putative NTD-NPC2 complex when there is cholesterol in NPC2 side,[19,21] which is called Texas model and it was suggested based on NTD domain only, generates significant structural crash when superimposed on the cryo-EM structure, indicating that possible involvement of some structural re-orientation or displacement of NTD to adapt the NPC2. The fitting of Texas complex into the current simulation structure after 1.0 μs in Figure 4 shows no more such crash and favorable interaction is possible between full length NPC1 and NPC2. Certainly, it would be interesting to keep track of this putative full length NPC1 + NPC2 structure with long time molecular dynamics in the presence of cholesterol, which is underway.

**Figure 4.**
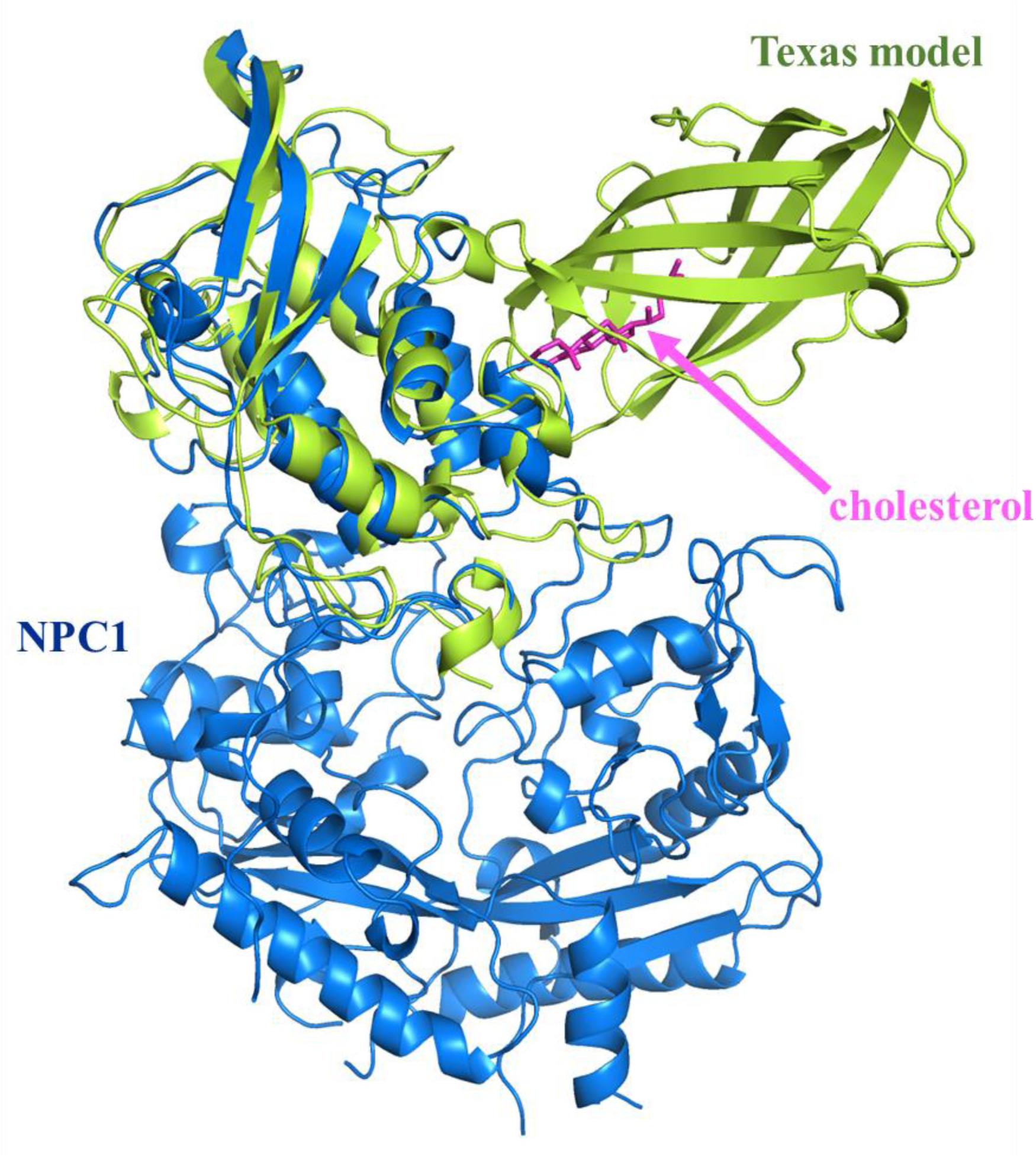
Fitting of the Texas structure (lime) onto the structure obtained from current simulation (blue).

As for R518W mutation simulation, we generated six independent trajectories for 200 ns each. Basically, no noticeable changes are observed except in one trajectory where the NTD is displaced toward the MLD side, which is opposite direction to the wild type simulation. Possibly, this causes less favorable interaction with NPC2 and could be the signature of the experimentally observed cholesterol independent weak interaction of NPC2 with mutated NPC1.[10,12] In fact, the overlap of MLD from the last frame of this trajectory over the MLD of X-ray structure (PDB ID: 5KWY)[16] generates structural crash in interface between NPC2 and NTD, more specifically residue A34∼K38 of NTD and residue G80∼P84 of NPC2 (Figure S2). Another possible cause of weaken interaction is the structural change of NPC2 binding MLD loops upon the mutation. It is reported that the seven residues (Q421, Y423, P424, D502, F503, F504, and Y506) in two protruding MLD loops are involved in interaction with NPC2. To understand this possibility, we analyzed the loop structure between wild and mutated trajectories but we was not able to correlate the loop structural difference with NPC2 binding affinity between them. Note that the residues D502 is E502 in our simulation since we have used human NPC1 unlike NPC1 X-ray structure, which is for bovine. Therefore, we believe the reduced binding affinity of mutated R518W is due to the displacement of NTD toward the MLD, leading to less favorable NTD-NPC2 interaction compare to a state where the interface is water exposed. This can be understood by noting that the interactions in this region is mostly between hydrophobic residues (or charged residues) from NTD and hydrophobic residues from NPC2.

We also performed the molecular dynamics simulation with cholesterol loaded in NTD. The seven independent simulations to 200 ns show no distinctive titling motion or distance change between domains. There was no noticeable structural differences between any two different trajectories either. Therefore we selected one of the trajectory and proceeded the simulation to 2.0 μs. Then, we observed change of NTD orientation. The backbone RMSD using starting structure as a reference was plotted in Figure 5 as a function of time. We observed that the trajectory quickly makes conduit-like channel as a result of this re-orientation, including the cholesterol containing NTD as a part of this channel, that was mentioned above. As one can see from Figure 6, which is the same angle profile as Figure 2, its relative orientation between domains are different from structure with no cholesterol at least within our simulation time scale. We have selected a structure near 1.8 μs which has RMSD of 9.0 Å and presented the identified tunnel in Figure 7. The tunnel was constructed using MoleOnline[44] and imported into PyMOL. It is somewhat surprising that this long tunnel is keep surviving even with large structural fluctuation. This tunnel might act as a channel for cholesterol transport from NTD to SSD much like the experimentally observed channel in NCR1.[26] Of course, the simulation is not reached to the fully equilibrated state. Hence, the tunnel we have observed should be understood as a snapshot of dynamically fluctuating system and its shape might be not ideal. However, this simulation shows the trajectory can form relatively stable tunnel, which was not observed in trajectory with no cholesterol on NTD, that can facilitate the transport of cholesterol and there is onset of NTD alignment to this channel when there is cholesterol on NTD, possibly for proper transport of cholesterol from NTD all the way to SSD. To observe the behavior of the mutated NPC1 once the cholesterol is transferred to NTD from NPC2, which might be relatively easy after binding of NPC2 to MLD loops, we performed the same simulations with cholesterol in NTD for R518W mutation. The number of total independent simulation trajectory was nine and the simulation was run up to 200 ns each for each of eight trajectories and 1.6 μs for one trajectory. Among these nine trajectories two of them shows similar behavior, i.e. formation of tunnel or tunnel-like structure as shown in Figure 8. We also examined the possibility of blocking of the tunnel due to the R518W mutation since it is located near the tunnel. But those tunnel blocking was not observed in all simulations. These observations are in agreement with experimental findings that the reduced cholesterol trafficking of R518W NPC1 is mainly due to the functional inability instead of misfolding as stated above.

**Figure 5.**
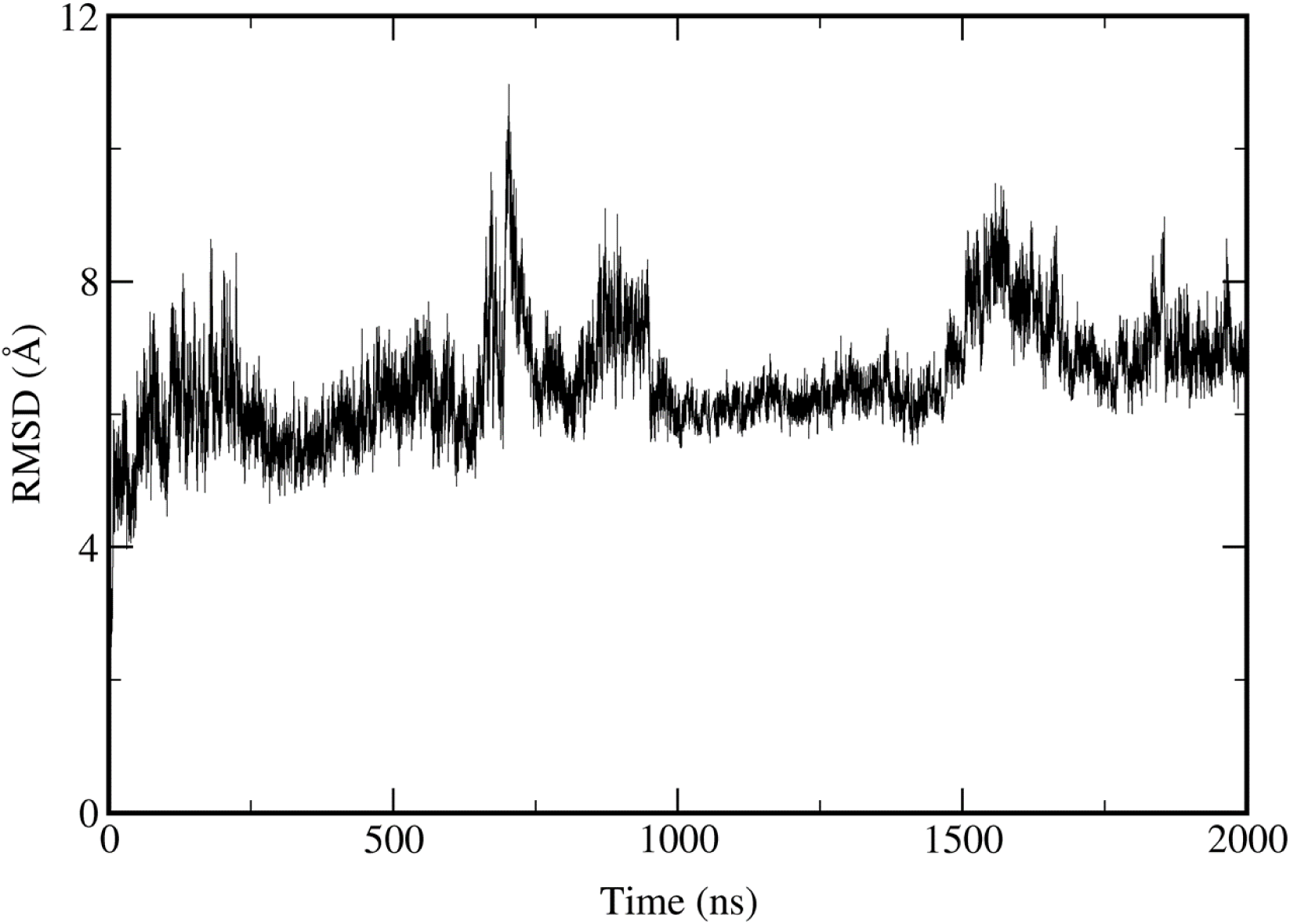
The RMSD time profile of wild type NPC1 simulation with cholesterol in NTD.

**Figure 6.**
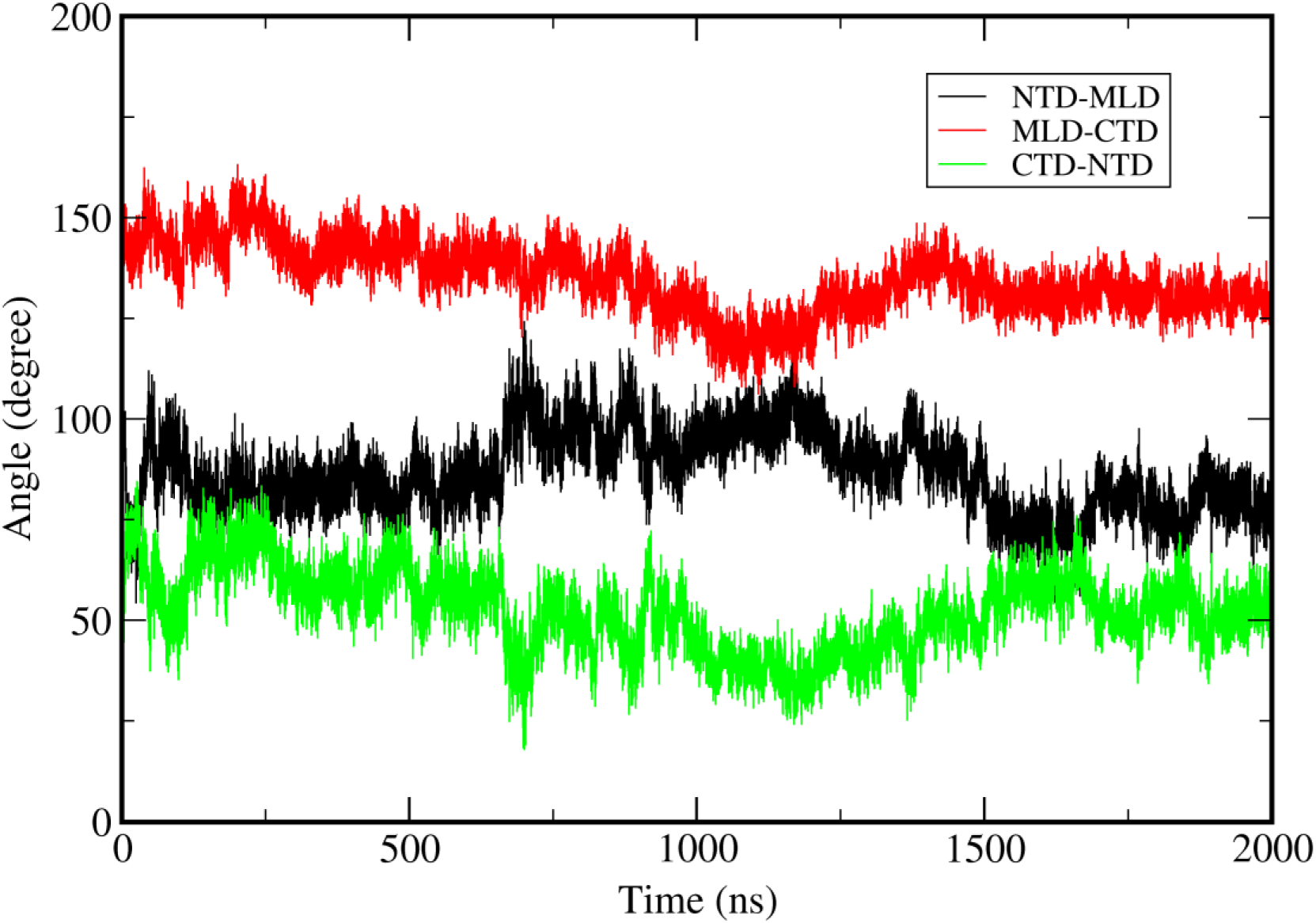
The change of angles as a function of time for a wild type trajectory with cholesterol in NTD. The angles here are obtained using the same method as in Figure 2.

**Figure 7.**
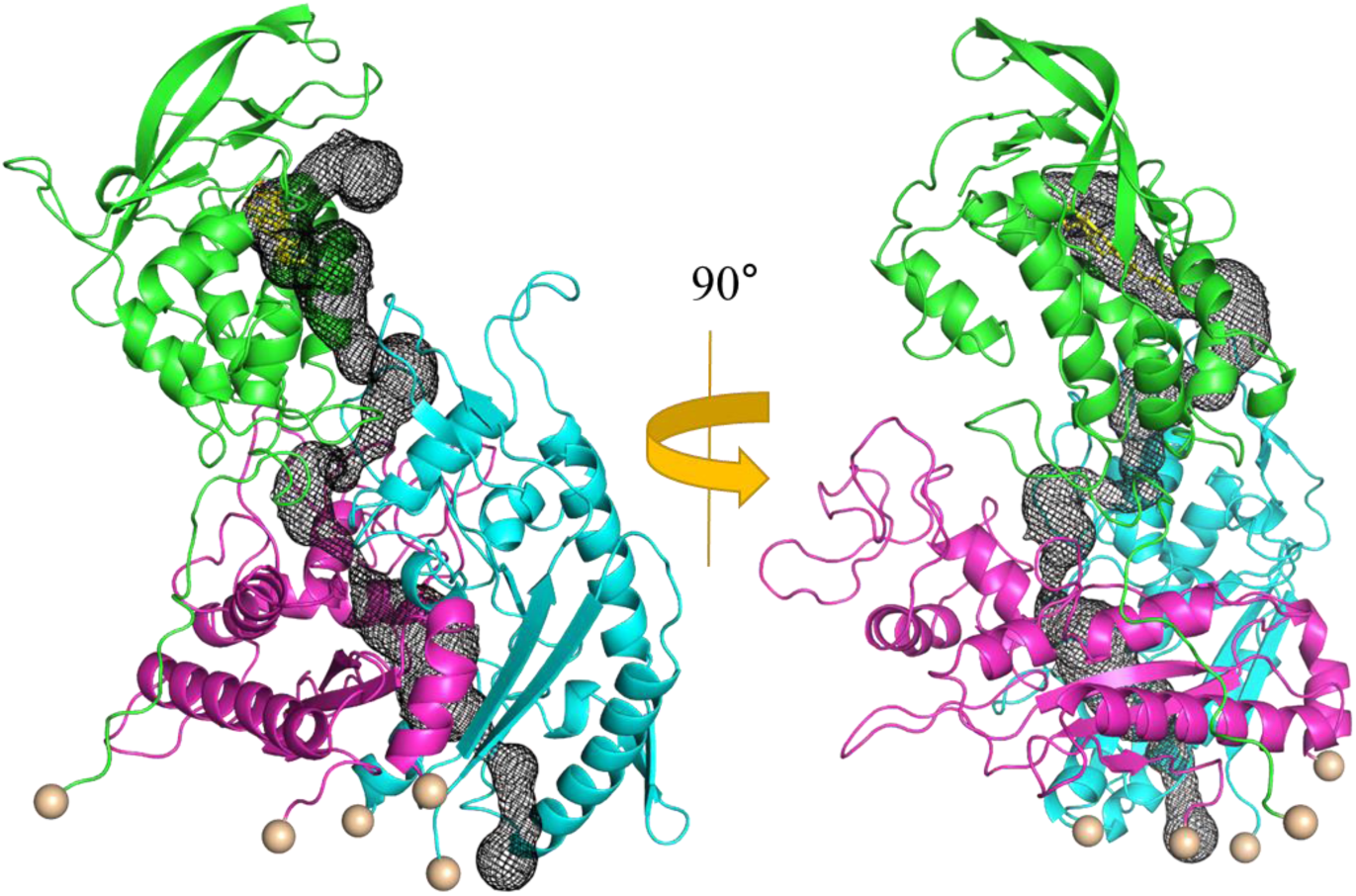
The tunnel obtained from wild type cholesterol containing NTD simulation. The tunnel is identified with MoleOnline[44] and represented as black mesh.

**Figure 8.**
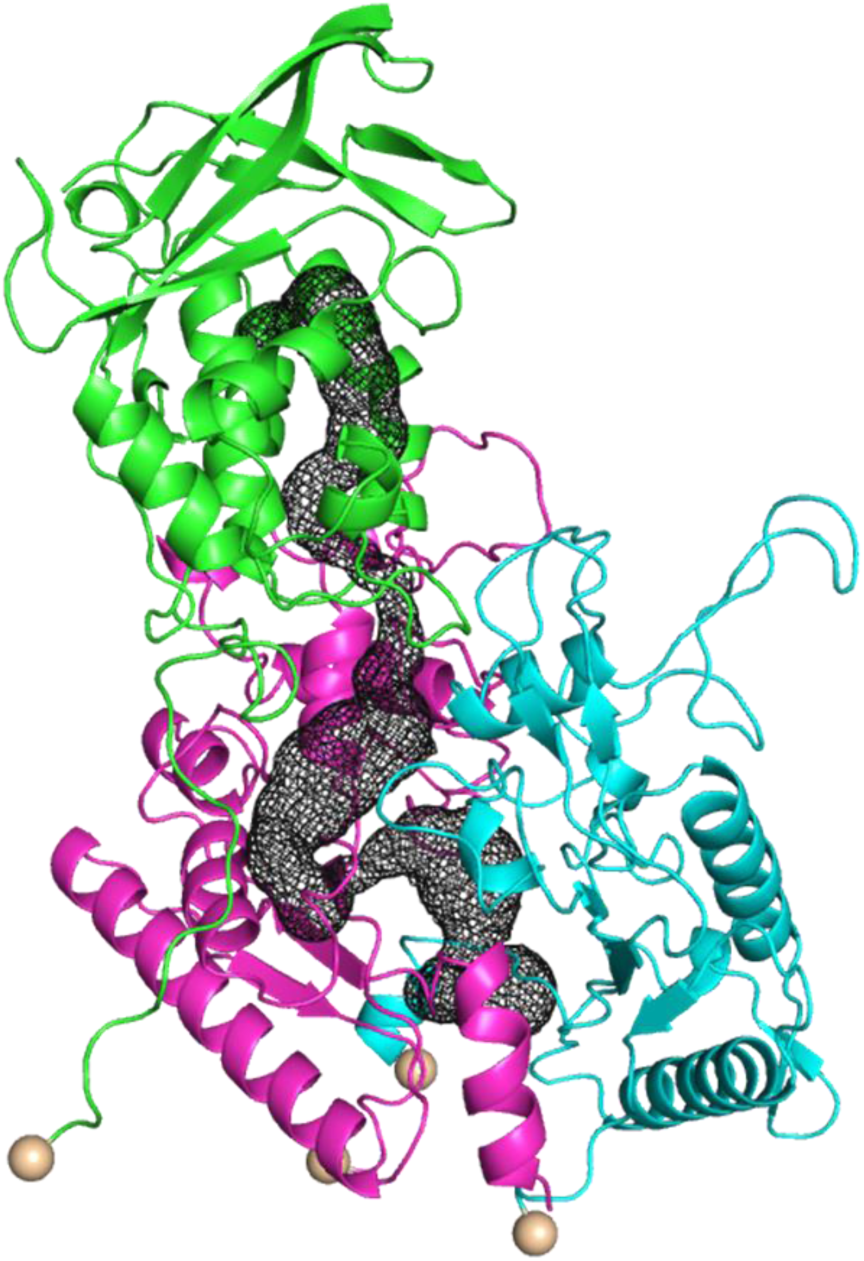
The same Figure as Figure 7 with R518W mutation with cholesterol in NTD.

## Conclusion

Based on the mutational study here, it seems that the reduced binding affinity NPC2 to R518W mutated NPC1 is due to unfavorable orientation of NTD for NPC2 binding, thereby decreased cholesterol transfer activity from NPC2 to NTD. On the other hands, it seems there is no difficulty of proper alignment of NTD to cholesterol transporting channel after the cholesterol is transferred to NTD from NPC2, supporting the stepwise process for cholesterol transport from NPC2 to SSD in lysosomal membrane. Before the transfer of cholesterol from NPC2 to NTD, the NTD has to have proper pose for favorable interaction with NPC2 as indicated by titling of NTD from current simulation and previous simulation.[23] The structure of full length NPC1 + NPC2 (with cholesterol) is not observed and it would be interesting to model this complex starting from putative structure presented in this work. The Texas model is based on NTD and NPC2 interaction and there is possibility that their interaction structure might be very different under the presence of full length NPC1. Right after the cholesterol transfer from NPC2 to NTD, the alignment of cholesterol in NTD must be directed more or less to NPC2 cholesterol leaving region. It is believed that the next step is the re-orientation of NTD starting from this conformation toward the long channel that leads to SSD, which could happen between neighboring NPC1. This process of leaving NPC2 from NTD and NTD re-orientation could happen independently or they could be connected. In either case, it seems reasonable to assume that from our simulations there is change of NTD orientation toward the favorable formation of tunnel that can facilitate the release of cholesterol from NTD to the tunnel for further process. This behavior was not observed previously.

We note that the cholesterol transfer efficiency from NPC2 to NPC1 is reduced by 90 % by deletion of NTD.[10] Therefore, the role of NTD must be crucial for proper cholesterol transport through NPC1. With current study, we demonstrated the importance of NTD as a dynamic domain depending on the existence of cholesterol on it along with the mutation R518W.

Clearly, there is limitation of current simulations since our model include lysosomal lumenal domains only instead of full length NPC1 including the membrane and cytoplasmic loops. Therefore the possible consorted motion between domains via the TMD and cytoplasmic loops is not fully reflected in current study. Moreover, the current simulation is too short to access the equilibrium state and the simulation results presented here should be understood as ‘dynamical signature’ of corresponding behaviors instead of final equilibrium property. Nevertheless, the current study provides insights into the structure function relation of NPC1, especially the shift of NTD orientation depending on the stage of cholesterol transport in connection to the NPC2 and effect of these motions or NPC1 lumenal domain conformations by R518W mutation. We believe the current study can enhance our understanding of cholesterol absorption/re-absorption process via NPC1 with atomic detail.

## Abbreviations

NPC: Niemann-Pick type C
NPC1: Niemann-Pick type C1
NPC2: Niemann-Pick type C2
MD: Molecular dynamics
NTD: N-terminal domain
MLD: Middle lumenal domain
CTD: C-terminal luminal domain
TMD: Transmembrane domain

## Acknowledgements

This research was supported by the National Research Foundation of Korea (NRF) grant funded by the Korean government (NRF-2018R1D1A1B07040808) to HJY. SJ acknowledges National Research Foundation of Korea (NRF) 2017M3D9A1073784.

## Author contributions

H.-J.Y., H.H.L. and S.J. conceived and designed the experiments. H.J performed computations. All authors analyzed the data, and wrote and edited the manuscript.

