## Supplementary material for "Signature of N-terminal domain (NTD) structural re-orientation in NPC1 for proper alignment of cholesterol transport: Molecular dynamics study with mutation": supplimentary

Hye-Jin Yoon[1†], Hyunah Jeong[2†], Hyung Ho Lee[1*], and Soonmin Jang[2*]
[1]Department of Chemistry, Seoul National University, Seoul, Republic of Korea,
[2]Department of Chemistry, Sejong University, Seoul, Republic of Korea

[†] These authors have equal contribution

### This PDF file includes:

Supplementary Figures. S1 to S2
Caption for Supplementary Movie

**List of Supplementary Figures**

Figure S1. The three helixes we selected for angle calculation between domains during the simulation. The residue number of each helix are Ser99~Thr112, Glu575~Asn593, and Phe1051~Met1069 in NTD, CTD, and MLD domains respectively.
Figure S2. The overlap of structure obtained from mutation simulation over the X-ray structure of NPC2 that is in binding with MLD. It generates some structural crash near the interface between NPC2 and NTD. The detailed structure of interface is shown along with residues.

Supplementary Movie in separate file. The movie of wild type NPC1 with no cholesterol on NTD starting from the initial structure. The length of the simulation in movie is 2.0 μs.

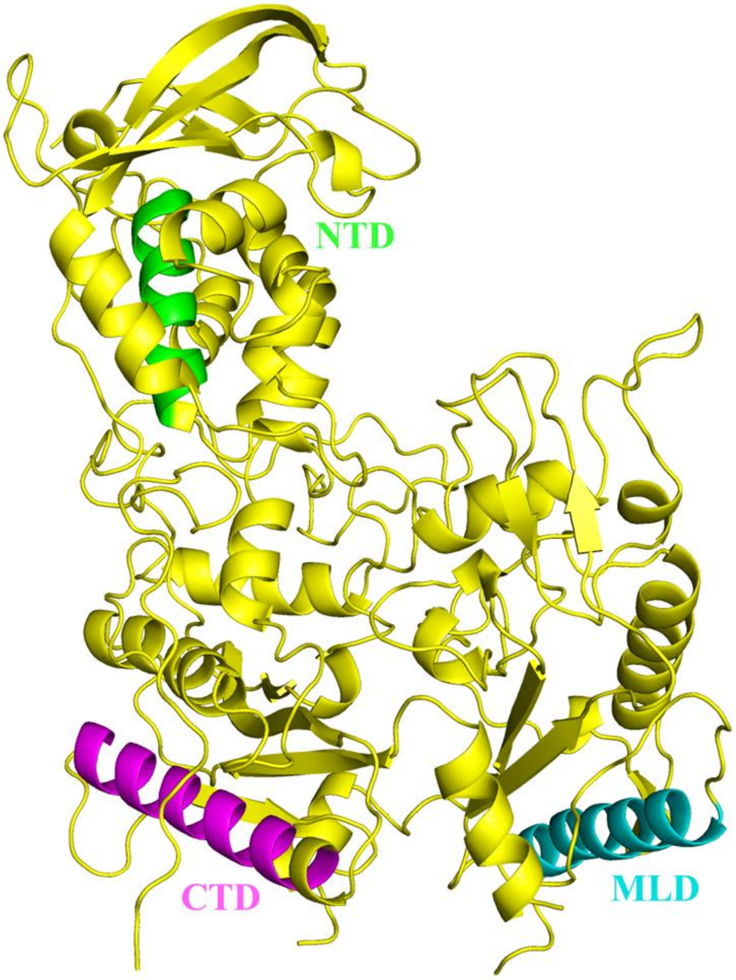


Figure S1

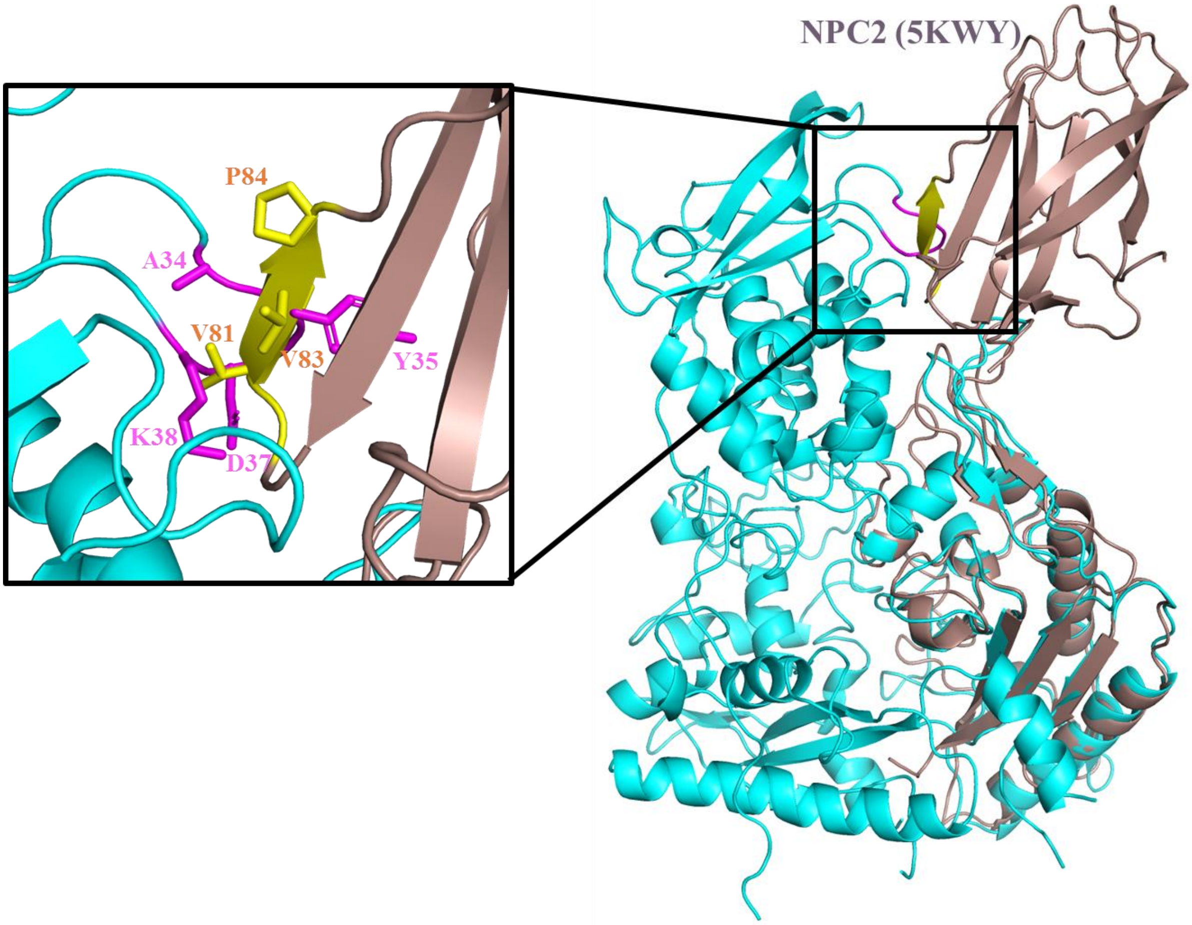


**Figure S2**
